# MRI tracking reveals selective accumulation of stem cell-derived magneto-extracellular vesicles in sites of injury

**DOI:** 10.1101/2019.12.22.885764

**Authors:** Zheng Han, Senquan Liu, Yigang Pei, Zheng Ding, Yuguo Li, Robert G. Weiss, Peter C.M. van Zijl, Jeff W.M. Bulte, Linzhao Cheng, Guanshu Liu

**Affiliations:** Russell H. Morgan Department of Radiology, Johns Hopkins University School of Medicine, Baltimore, MD, USA; F.M. Kirby Research Center, Kennedy Krieger Institute, Baltimore, MD, USA; Cellular Imaging Section and Vascular Biology Program, Institute for Cell Engineering, Johns Hopkins University School of Medicine, Baltimore, MD, USA; Department of Medicine, Johns Hopkins University School of Medicine, Baltimore, MD, USA; Department of Radiology, Xiangya Hospital, Central South University, Changsha, Hunan, China; Division of Cardiology, Department of Medicine, Johns Hopkins University School of Medicine, Baltimore, MD, USA

**Keywords:** MRI, magnetic nanoparticle, extracellular vesicle, stem cell, iPSC, acute kidney injury

## Abstract

Human stem-cell-derived extracellular vesicles (EVs) are currently being investigated for cell-free therapy in regenerative medicine applications, but their biodistribution and tropic properties for homing to injured tissues are largely unknown. Here, we labeled EVs with magnetic nanoparticles to create magneto-EVs that can be tracked by magnetic resonance imaging (MRI). Superparamagnetic iron oxide (SPIO) nanoparticles were coated with polyhistidine tags, which enabled purification of labeled EVs by efficiently removing unencapsulated SPIO particles in the solution. The biodistribution of systemically injected human induced pluripotent stem cell (iPSC)-derived magneto-EV was assessed in three different animal models of kidney injury and myocardial ischemia. Magneto-EVs were found to selectively home to the injury sites and conferred substantial protection in a kidney injury model. *In vivo* MRI tracking of magnetically labeled EVs represents a new powerful method to assess and quantify their whole-body distribution, which may help optimize further development of EV-based cell-free therapy.

## Introduction

Extracellular vesicles (EV) are small membranous blebs or vesicles released from nearly all cell types that function as important messengers and mediators for intercellular communication in a diverse range of biological processes(*1*). Depending on their biogenesis(*2*), EVs are typically classified into exosomes (50–150 nm in diameter) or microvesicles (50-500 nm in diameter). Exosomes are derived from specialized intracellular compartments, *i.e.*, endosomes or multi-vesicular bodies (MVBs), while microvesicles are shed directly from the plasma membrane. When reaching recipient cells that are either in the immediate vicinity or at a distance, EVs fuse with them to transmit cargo (*e.g.*, proteins, messenger RNA, microRNA, and lipids) and elicit biologic changes in recipient cells(*3*). In stem cell research, increasing evidence shows that EVs are essential for stem cells to protect or regenerate injured cells, presumably through an EV-mediated paracrine effect(*4, 5*). This has led to the use of stem cell-derived EVs alone for cell-free therapy(*6, 7*). Indeed, compared to parental mesenchymal stem cells (MSCs) and induced pluripotent stem cells (iPSCs), human stem cell-derived EVs are considered to be a safer and more effective regenerative medicine approach for treating many otherwise untreatable diseases such as kidney injury(*8-11*) and cardiac disease(*12, 13*). Despite the appeal and potential impact of EV-based therapies, there are currently no clinic-ready methods for noninvasively tracking their bio-distribution and uptake to guide, optimize, and monitor treatment.

Noninvasive *in vivo* imaging approaches are needed that can track the distribution and fate of administered EVs over time. Importantly, such methods could be used to assess the quantity of administered EVs in the organs to be treated, which may be predictive of a therapeutic response. Before this, EV imaging and tracking methods can serve as a guide to direct the development, optimization, and implementation of EV-based therapeutics. To date, considerable effort has been devoted to develop EV tracking methods using fluorescence imaging(*14, 15*), bioluminescence imaging(*16*), single-photon emission computed tomography (SPECT)(*17*), positron emission tomography (PET)(*18*), computed tomography (CT)(*19, 20*), magnetic resonance imaging (MRI)(*21, 22*), and magnetic particle imaging(*23*). Among them, MRI is an appealing imaging modality that is used widely in the clinic and has excellent soft-tissue contrast without using ionizing radiation. To the best of our knowledge, there are only two reports on *in vivo* MRI-based EV-tracking studies. In the study reported by Busato *et al*(*21*), superparamagnetic iron oxide (SPIO) particles were introduced into the parent cells (adipose stem cells) and the EVs secreted from these cells were then collected. While this approach doesn’t require a purification procedure to remove free SPIO, the labeling efficiency was unfortunately found to be low, making *in vivo* tracking of EVs challenging due to the inherently low sensitivity of MRI. Hu *et al.* used electroporation to load SPIOs into EVs and subsequently purified them by ultracentrifugation(*22*). While this approach provides high labeling, ultracentrifugation is not an efficient method to remove free naked SPIO nanoparticles and induces SPIO aggregation when pelleting. Hence, an efficient technology to obtain highly purified SPIO-labeled EVs is still an unmet need.

The aim of this study was to first develop a platform technology for preparing highly purified magnetically labeled EVs (magneto-EVs) and then to test *in vivo* MRI tracking of their whole-body distribution. We chose to use iPSC-derived EVs because iPSCs are more amenable to generate therapeutic EVs to protect or repair injury, which has been demonstrated in a few preclinical studies(*24*). We have recently shown that human iPSCs can be cultured infinitely in chemically defined medium (free of exogenous EVs from serum or other biological sources) and produce EVs markedly higher than other types of stem cells(*25*). Considering that stem cell derived EVs lack nuclei, iPSC-derived EVs are attractive without the concerns of genetic instability and tumorigenicity associated with the administration of whole iPSCs(*26, 27*). Moreover, individualized autologous iPSCs can be generated from a single blood draw or skin sample from the same patient to minimize immunogenicity and ethical concerns(*26*). Using different animal tissue injury models, we show here that the temporal-spatial distribution of magnetically labeled iPSC-derived EVs can be assessed quantitatively *in vivo* with MRI.

## Results

### Magnetic labeling of EVs and purification

To prepare “sticky” magnetic nanoparticles, we first synthesized surface-modified SPIO nanoparticles by conjugating them with hexa-histidine peptides (6×His) using the synthetic route shown in **Fig. 1A**. With His-tags on the surface, which was confirmed by Fourier-transform infrared spectroscopy (**Fig. S1**), SPIO nanoparticles can selectively bind to Ni^2+^ immobilized nitrilotriacetic acid (NTA) agarose resins (Ni-NTA) (**Fig. 1B)**. This complex has a distinct color (**Fig. S2)**, where the Ni-NTA column eluted with SPIO-COOH (no His tag) showed no color change whereas the one eluted with SPIO-His changed from light blue to brown.

**Figure 1.**
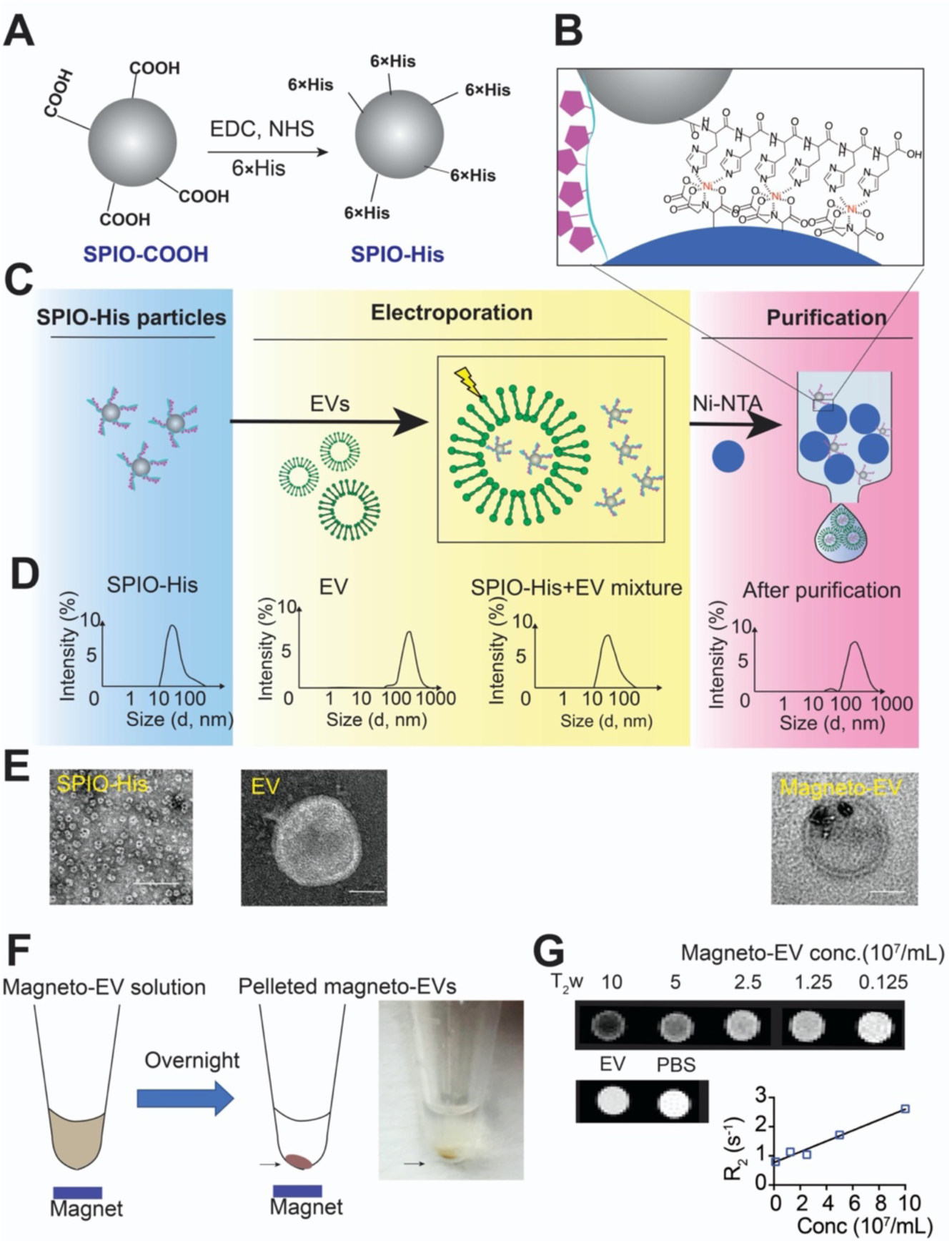
Preparation magneto-EV and characterization of purified magneto-EVs. **A.** Schematic illustration of the preparation of SPIO his-tag (SPIO-His), by conjugating hexahistidine (6×His-tag) polypeptide to the carboxyl groups of SPIO particles using EDC (1-Ethyl-3-(3-dimethylaminopropyl)-carbodiimide), and NHS (sulfo-N-hydroxysuccinimide) chemistry. **B.** As a result from the high affinity between the His-peptide and nickle ion, the SPIO-His particles bind to Ni^2+^ immobilized on beads (e.g., Ni-NTA resins) for further purification. **C**. Schematic illustration of the encapsulation of SPIO-His into EVs by electroporation and subsequent purification by removing unencapsulated SPIO-His from the elute using Ni-NTA affinity chromatography. **D.** Size distribution as measured by dynamic light scattering (DLS) for SPIO-His, EVs, SPIO-His/magneto-EV/EV mixtures after electroporation, and the final purified elute, respectively. **E**. TEM images of EV, SPIO-His and EV-SPIO, respectively. **F.** Concentration of magneto-EVs using a magnet. Eluted magneto-EV solution was placed on a magnet overnight to pellet magneto-EVs. The photograph shows the pelleted magneto-EV at the bottom of a microcentrifuge tube. **G**. T_2_-weighted (T_2_w) images of magneto-EVs at different concentrations, unlabeled EVs, and PBS. Mean R_2_ values of magneto-EVs are plotted with respect to their concentration, from which the r_2_ (relaxivity) was estimated.

The labeling and purification procedures of magneto-EVs are illustrated in **Fig. 1C**. In brief, the synthesized SPIO-His nanoparticles were first loaded into purified EVs by electroporation as described previously(*25*). The resulting solution containing a mixture of free SPIO-His and magneto-EVs was purified using a Ni-NTA column. Quantitative analysis of iron content in the solution pre- and post-elution revealed that the Ni-NTA column was able to remove 97.4% of unincorporated SPIO-His with minimal loss of EVs (∼5.4% as measured by nanoparticle tracking analysis). The average size was measured using dynamic light scattering (DLS) to be 43.9±16.5 nm, 248.2±107.2 nm, 45.6±15.9 nm, and 292.8±113.0 nm, for SPIO-His, EVs, unpurified magneto-EVs, and purified magneto-EVs, respectively (**Fig. 1D**). The size distribution of magneto-EVs closely resembled that of unlabeled EVs, indicating the efficient removal of unencapsulated SPIO-His particles. Labeling and purification were verified by transmission electron microscopy (TEM) (**Fig. 1E**), which showed no unencapsulated SPIO-His particles in the purified solution. TEM demonstrated that many EVs contained multiple SPIO-His particles. Moreover, the incorporation of magnetic particles allowed enrichment of magneto-EVs using a magnetic force (**Fig. 1F**).

*In vitro* MRI confirmed the hypointense contrast of magneto-EVs (**Fig. 1G**). At 9.4 T and 37 °C, the R_2_ enhancement was determined to be 1.83 s^-1^ per 10^8^/mL EVs, which is equivalent to an r_2_ relaxivity of 1.1×10^10^ s^-1^mM^-1^ per EV (10^8^/mL EVs = 16.7 pM) or 659 s^-1^mM^-1^ Fe (10^8^/mL EVs contain 155 ng/mL Fe as measured by ICP-OES). From this, we estimated the *in vivo* detection limit to be approximately 8.76×10^7^ EVs/mL EVs kidney tissue (having an inherent R_2_ of 32.03 s^-1^ at 3T(*28*)), assuming a 5% MRI signal change.

### In vivo MRI of iPSC-EV uptake in an injured kidney model

We first assessed the distribution of magneto-EVs in an lipopolysaccharides (LPS)-induced acute kidney injury (AKI) model, a well-established rodent model associated with severe systemic inflammation and irreversible kidney damage within 24-48 hours(*29, 30*). As SPIO nanoparticles strongly distort the local magnetic field and result in a much quicker T_2_* decay of protons in nearby water molecules(*31, 32*), the dynamic uptake of SPIO-containing magneto-EVs could be readily detected using T_2_*-weighted (T_2_*w) MRI (**Supplementary video S1**) in which magneto-EVs appear as hypointense spots. As shown in **Fig. 2A**, either SPIO-His particles or magneto-EVs were *i.v.* injected into uninjured control or LPS-AKI mice and the MRI signal was monitored dynamically for 30 minutes after injection. The 30-minute window was chosen because the blood half-life of EVs has been reported to be 2-4 minutes only(*33*).

**Figure 2.**
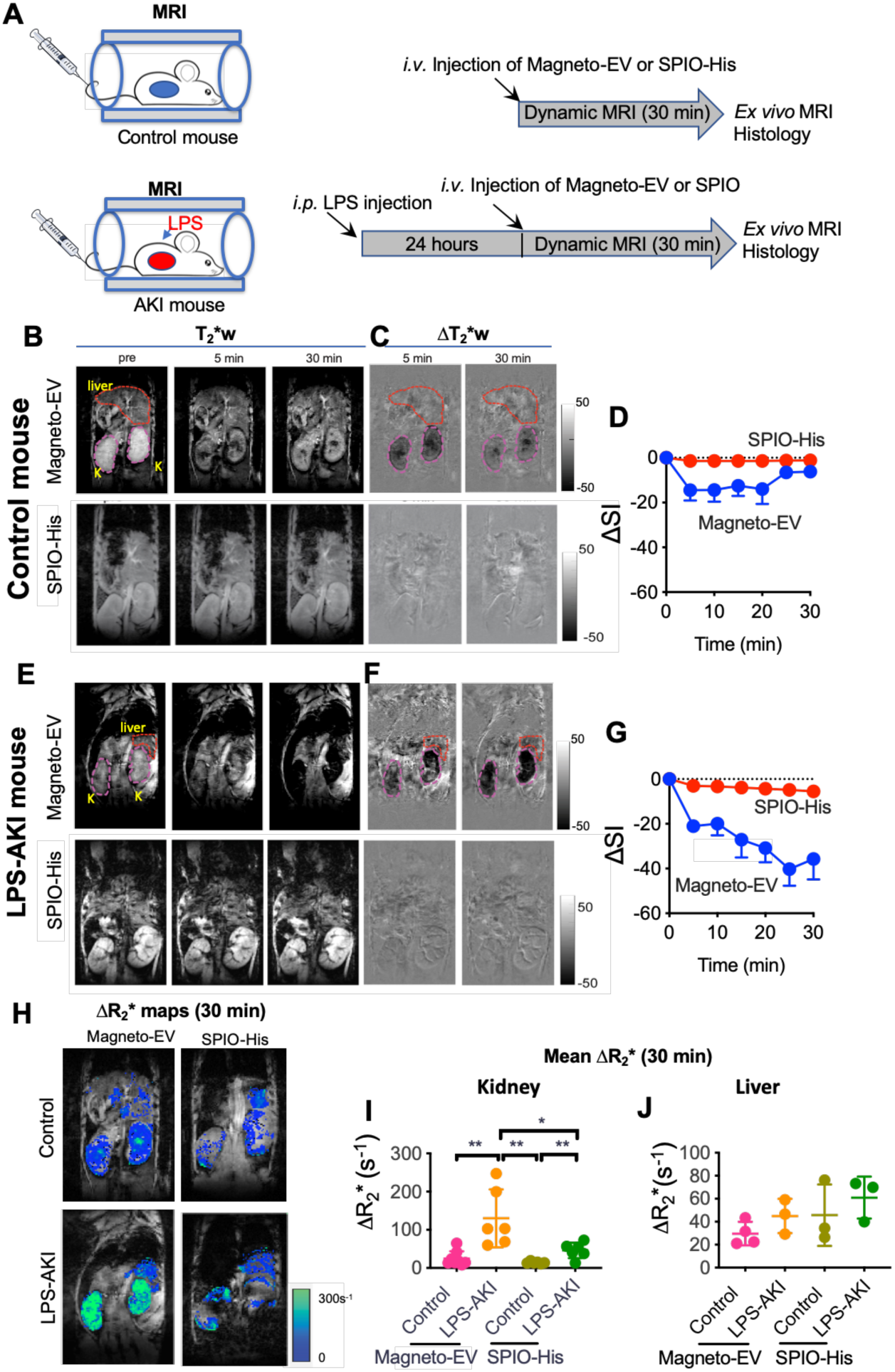
MRI detection of the uptake of magneto-EV in control and injured kidneys in the LPS-AKI model. **A**. Timeline of preparation of animal model and injection of EVs. **B**. T_2_*-weighted (T_2_*w) images in a representative normal control mouse before and at 5 and 30 min after *i.v.* injection of magneto-EVs and SPIO-His, respectively. **C**. Corresponding contrast enhancement maps, defined as ΔT_2_*w=T_2_*w (post)-T_2_*w (pre). **D.** Mean dynamic signal change in the control kidneys (n=6). **E**. T_2_*-weighted (T_2_*w) images in a representative LPS-AKI mouse before and at 5 and 30 min after the *i.v.* injection of magneto-EVs and SPIO-His, respectively. **F**. Corresponding contrast enhancement maps. **G.** Mean dynamic signal change in the injured kidneys (n=6). **H**. ΔR_2_*(1/T_E_ × ln(SI^post^/SI^pre^)) maps at 30 min after injection of magneto-EVs. **I.** Quantitative comparison of mean kidney ΔR_2_* values at 30 min between different groups. *, *P*<0.05, ns: not significant, n=6, unpaired two-tailed Student’s t-test. **K.** Quantitative comparison of mean liver ΔR_2_* values at 30 min between different groups. *, *P*<0.05, ns: not significant, n=6, unpaired two-tailed Student’s t-test.

For uninjured control mice, the MRI signal intensity in the kidney exhibited a rapid decrease immediately after *i.v.* injection of magneto-EVs and then remained stable between the 5 to 20 min, finally recovered to baseline at 20 -30 min (**Figs. 2B-D**). In contrast, when SPIO-His nanoparticles were injected at the same iron dose, no detectable signal change in the kidney could be observed. In the LPS-AKI mice (**Figs. 2E-G**), a substantially different uptake pattern was observed in animals injected with magneto-EVs, with the MRI signal continuing to decrease for ∼25 min when it reached a plateau, indicating a continuous uptake in the injured kidneys. At 30 min after magneto-EV injection, the injured kidneys nearly lost all their signal, indicating a high uptake of magneto-EVs. LPS-AKI mice injected with SPIO-His nanoparticles alone showed only negligible MRI signal changes in the kidney, similar to the mice in the uninjured control group injected with SPIO-His particles.

The amount of magneto-EVs or SPIO-His particles accumulated in the kidney was then quantitatively estimated using the changes of R_2_*signal in the kidney, defined as ΔR_2_* =1/T_2_*(post)-1/T_2_*(pre) =1/T_E_ × ln (SI^post^/SI^pre^), a commonly used metric in SPIO-enhanced MRI(*34*). Snapshot ΔR_2_* contrast enhancement maps and changes at 30 minutes after magneto-EV injection are shown in **Fig. 2H**. A higher amount of magneto-EVs accumulated in injured vs. uninjured control kidneys. Weaker ΔR_2_* contrast enhancement was seen in both groups of kidneys injected with SPIO-His nanoparticles. A quantitative comparison of kidney ROI values revealed a significantly higher ΔR_2_* for the LPS-AKI group compared to uninjured controls (130.1 vs 24.69 s^-1^, *P*=0.0024) (**Fig. 2I**). There was also a significant difference between the LPS-AKI mice injected with magneto-EVs vs. SPIO-His particles (130.1 vs. 46.0 s^-1^, *P*=0.0254). The ΔR_2_* in the LPS-AKI group after SPIO-His injection was also higher than that in the normal control group (46.0 vs, 14.3 s^-1^, *P*=0.0035). In addition, we calculated the dynamic ΔR_2_* values in the kidney for each group (**Fig. S3B**), confirming the differential kidney uptake dynamics for LPS-AKI mice injected with magneto-EVs versus all other groups.

To determine whether magneto-EVs can be taken up by the liver non-specifically, as is the case for most *i.v.-*injected SPIO nanoparticles(*35*), we further analyzed liver contrast-enhancement in both LPS-AKI and uninjured control mice (**Fig. S3A**). As shown in **Fig. 2J**, none of the groups showed significantly different ΔR_2_* values at 30 min after injection of magneto-EVs or SPIO-His particles. Similar dynamic liver uptake patterns were observed among these groups (**Fig. S3C**), suggesting that there is negligible non-specific liver uptake of iPSC-EVs.

### High-resolution ex vivo MRI of iPSC-EV uptake in an injured kidney model

To study the biodistribution of magneto-EVs and naked SPIO-His particles in kidneys in more anatomic detail, we performed high resolution three-dimensional (3D) *ex vivo* MRI on fixed kidney samples that were excised at 30 min after magneto-EV or SPIO-His injection. **Fig.3A** shows representative kidney T_2_*w images of LPS-AKI mice injected with magneto-EVs, SPIO-His particles, and saline, respectively. Only the kidneys from mice receiving magneto-EVs demonstrated a high number of hypointense spots and streaks dispersed throughout the renal cortex (**Fig. 3A, left**). In contrast, kidneys of mice injected with SPIO-His particles showed far fewer black spots (**Fig. 3A, middle**), similar to the kidney without SPIO-His injection (**Fig. 3A, right**). The presence of black hypointensities in the last two groups may be due to the LPS-induced hemorrhage (H&E stain, **Fig. S4**). 3D reconstruction of hypointense voxels revealed that magneto-EVs distributed throughout the whole kidney, with preferential accumulation in the cortex (**Fig. 3B**, full 3D visualization shown in **supplementary video 3)**. Quantitative analysis showed that approximately 28.1% of the cortex of AKI mice injected with magneto-EVs contained hypointensities (**Fig. 3C**), significantly higher than those injected with SPIO-His particles alone (12.5%, *P*<0.0001) or saline (11%, *P*<0.0001). Accumulation of magneto-EVs in the cortex was confirmed by Prussian blue staining for iron (**Fig. 3D**), where the tissue distribution of magneto-EVs showed a good agreement with that seen on *ex vivo* MRI. Furthermore, immunostaining for VCAM-1, a vascular inflammation marker, showed that LPS-AKI kidneys exhibited higher VCAM-1 expression in the cortex compared to the medulla (**Fig. 3E**). Prussian blue-positive iron also co-localized with dilated proximal tubules as shown on Periodic acid–Schiff (PAS) staining (**Fig. 3F**), with the tubular damage being a hallmark of the LPS-AKI model(*36*). In contrast, kidneys from control mice injected with either magneto-EVs or SPIO-His particles exhibited fewer hypointense areas, and those hypointense areas on T_2_*w images (**Fig. 3G**) correlated well with the negative Prussian blue staining (**Fig. 3H**).

**Figure 3.**
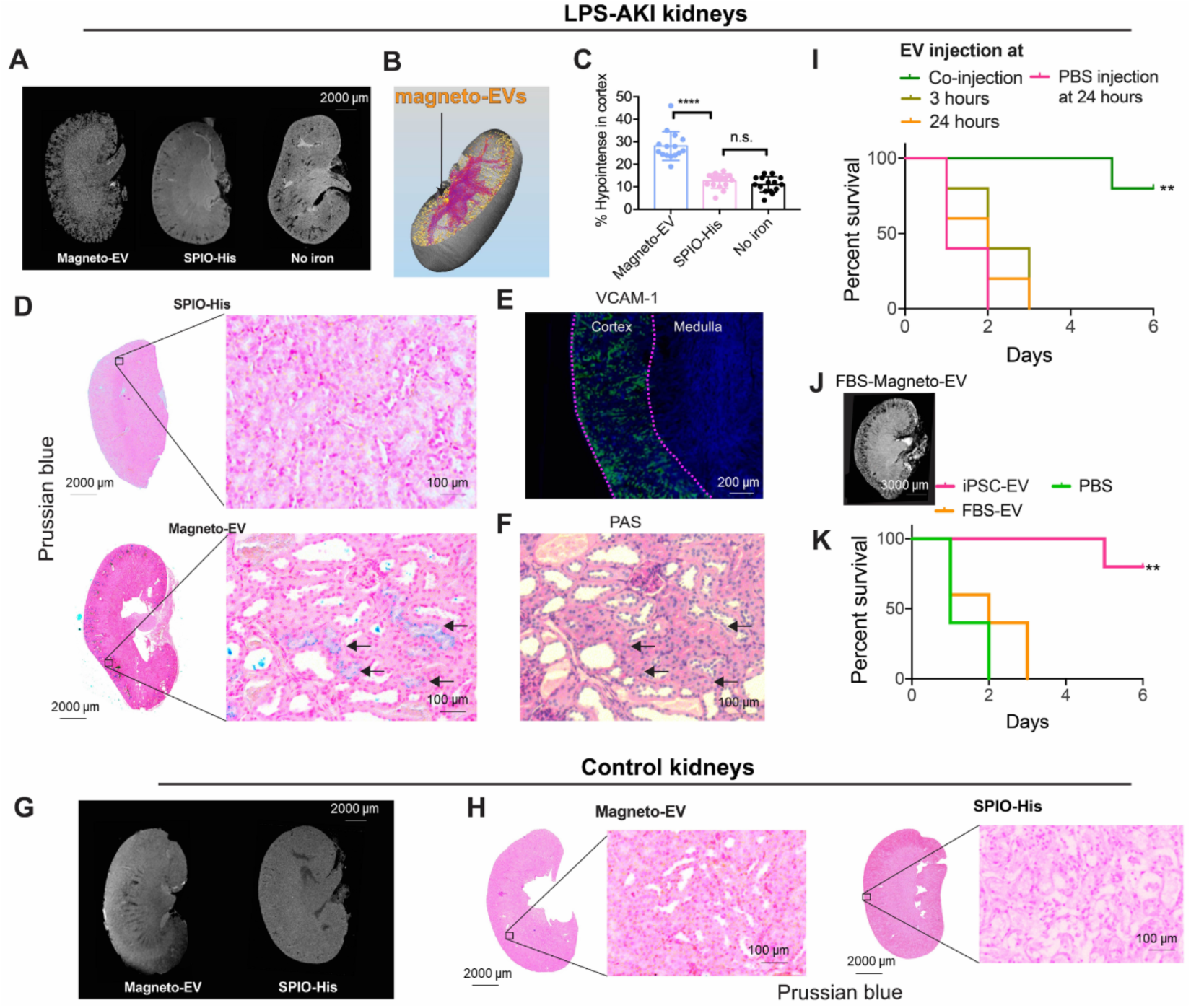
Biodistribution of magneto-EVs in the injured kidney and their therapeutic effect in the LPS-AKI model. **A**. Different biodistribution patterns of magneto-EVs (left) and SPIO-His (middle) in representative LPS-AKI kidneys, as revealed by *ex vivo* high-resolution MRI. Image of a representative non-injected kidney on the right is shown as a control reference. **B**. 3D reconstruction of a representative kidney showing the biodistribution of magneto-EVs (gold-colored dots). Blood vessels are shown in purple color (see also Supplementary Video 3). **C**. Quantitative comparison of the relative hypointense areas (%) in the cortex of AKI mice injected with magneto-EVs, SPIO-His, or without injection (****: *P*<0.0001, unpaired Student’s t-test, n=15). **D**. Prussian blue stains showing the distribution of magneto-EVs and SPIO in the injured kidney (Left:whole kidney; Right: zoom-in; Blue=SPIO; Red=nucleus). **E**. VCAM-1 staining of a representative kidney showing extensive inflammation (green) occurring in the cortex. Tissue was counterstained with DAPI (blue). **F**. Periodic acid–Schiff (PAS) staining of the section corresponding to the Prussian blue stain on the left. **G**. Biodistribution of magneto-EVs (left) and SPIO-His (right) in a representative normal control kidney. **H**. Prussian blue stains showing the distribution of magneto-EVs and SPIO-His in uninjured kidney (Left: whole kidney; Right: zoom-in; Blue=SPIO; Red=nucleus). **I.** Survival curves of AKI mice treated with 2×10^9^ iPSC-EVs at different time points (0, 3, and 24 hours) and vehicle control (PBS). A significant treatment response can be observed at 0 hours, i.e. co-injection of LPS and iPSC-EV (**: *P*=0.0023 vs. PBS, log-rank test). **J**. *Ex vivo* MR images of a representative LPS-AKI kidney 30 min after injection 2×10^9^ FBS-derived magneto-EVs. **K.** Survival curves of LPS-AKI mice co-injected with iPSC-EV, FBS-EV, or PBS, respectively. **: *P*=0.0023, iPSC-EV vs. PBS; *P*=0.0025, iPSC-EV vs. FBS-EV; log-rank test.

The accumulation of iPSC-derived EVs in the injury sites resulted in an observable therapeutic effect. A single dose of 2×10^9^ EVs was injected at 0 (LPS co-injection), 3, or 24 h after LPS injection (**Fig. 3I**). Compared to vehicle control (PBS), only the magneto-EV/LPS co-injection resulted in a statistically significant improvement of survival rate (n=5, *P*=0.0023, unpaired two-tailed Student’s t-test). The lack of improvement of survival when magneto-EVs were injected at 3 and 24 hours is likely due to the rapid progression of kidney injury in the LPS-AKI model. Indeed, without treatment, the animal loss at 24 and 48 hours was 50 and 100%, respectively. To assess whether the accumulation of magneto-EVs in the injured kidney and their subsequent protective effect is attributed to their stem cell origin, we also isolated EVs from fetal bovine serum (FBS) and injected them into LPS-AKI mice using the same protocol as for iPSC-EVs. The *ex vivo* MRI results (**Fig. 3J**) showed that a high quantity of FBS-EVs accumulated in the cortex similarly to iPSC-EVs, suggesting that the accumulation of magneto-EVs in the injured sites may not be specific to their cell origin. However, despite their similar uptake pattern FBS-EVs did not generate protection against the LPS-induced kidney injury (**Fig. 3K**, *P*=0.2055), suggesting that the therapeutic effects are specific to the cell origin of the EVs.

### Biodistribution of iPSC-EVs in other experimental injury models

We further assessed the ability to track magneto-EVs in two other animal injury models. First, kidneys were subjected to a unilateral ischemia-reperfusion injury (IRI) (**Fig. 4A**). An acute injury in the right kidney was obtained by occluding the blood supply for 45 min, with the untreated left kidney as control. iPSC-derived magneto-EVs were administered *i.v*. at the same time when reperfusion started. As shown in **Figs. 4B,C**, a higher EV uptake was observed in the injured kidneys but not the contralateral ones (ΔR_2_*= 30.1 s^-1^ and 15.7 s^-1^, *P*=0.04, two-tailed paired Student’s t-test, n=4). The biodistribution pattern of magneto-EVs homing to the injury site was distinct from the AKI mice induced by LPS injection. Both the *ex vivo* MRI (**Fig. 4E**) and histology (**Fig. 4F**) showed more magneto-EVs accumulating in the medulla than the cortex, representing the difference in the primary site of injury between these two experimental models. We then applied our technology to study the distribution of *i.v.*-injected magneto-EVs in an IR-injured mouse heart, which was induced through the ligation of the left anterior descending (LAD) coronary artery for 35 min, followed by reperfusion(*37*) (**Figs. 5A,B**). The MRI results revealed that magneto-EVs accumulated selectively at the injury sites (**Figs. 5C,D**). The distribution of EVs followed an alignment with the injured myocardium (**Fig. 5E**), which was confirmed by high resolution *ex vivo* MRI (**Fig. 5F**) and Prussian blue staining (**Fig. 5G**).

**Figure 4.**
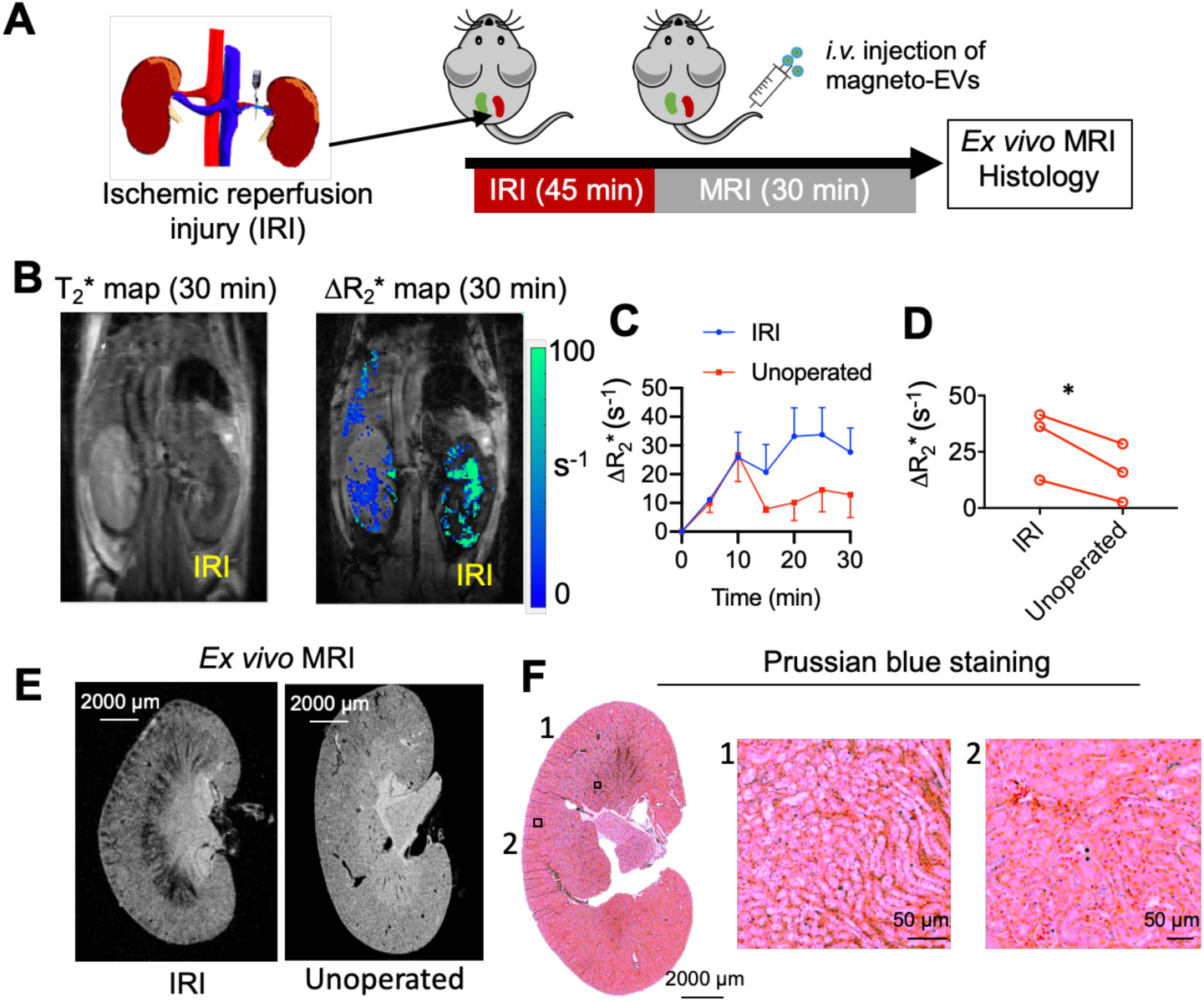
MRI tracking of *i.v.* administered magneto-EVs in the IRI-AKI model. **A**. Schematic illustration of the experimental IRI-AKI model and MRI acquisition. **B.** Representative *in vivo* T_2_* image and ΔR_2_* map at 30 min after EV injection. **C**. Dynamic ΔR_2_* MRI signal changes in IRI and non-operated control kidneys (n=3 in each group). **D**. Comparison of ΔR_2_* (30 min) values in IRI and control kidney for each mouse (*: *P*=0.04, two-tailed paired Student’s t-test, n=3). **E**. *Ex vivo* high-resolution T_2_*w MR image of a representative IRI kidney and unoperated kidney. **F**. Corresponding Prussian blue staining (Left: whole kidney; Right: zoom-in; Blue=SPIO; Red= nucleus.

**Figure 5.**
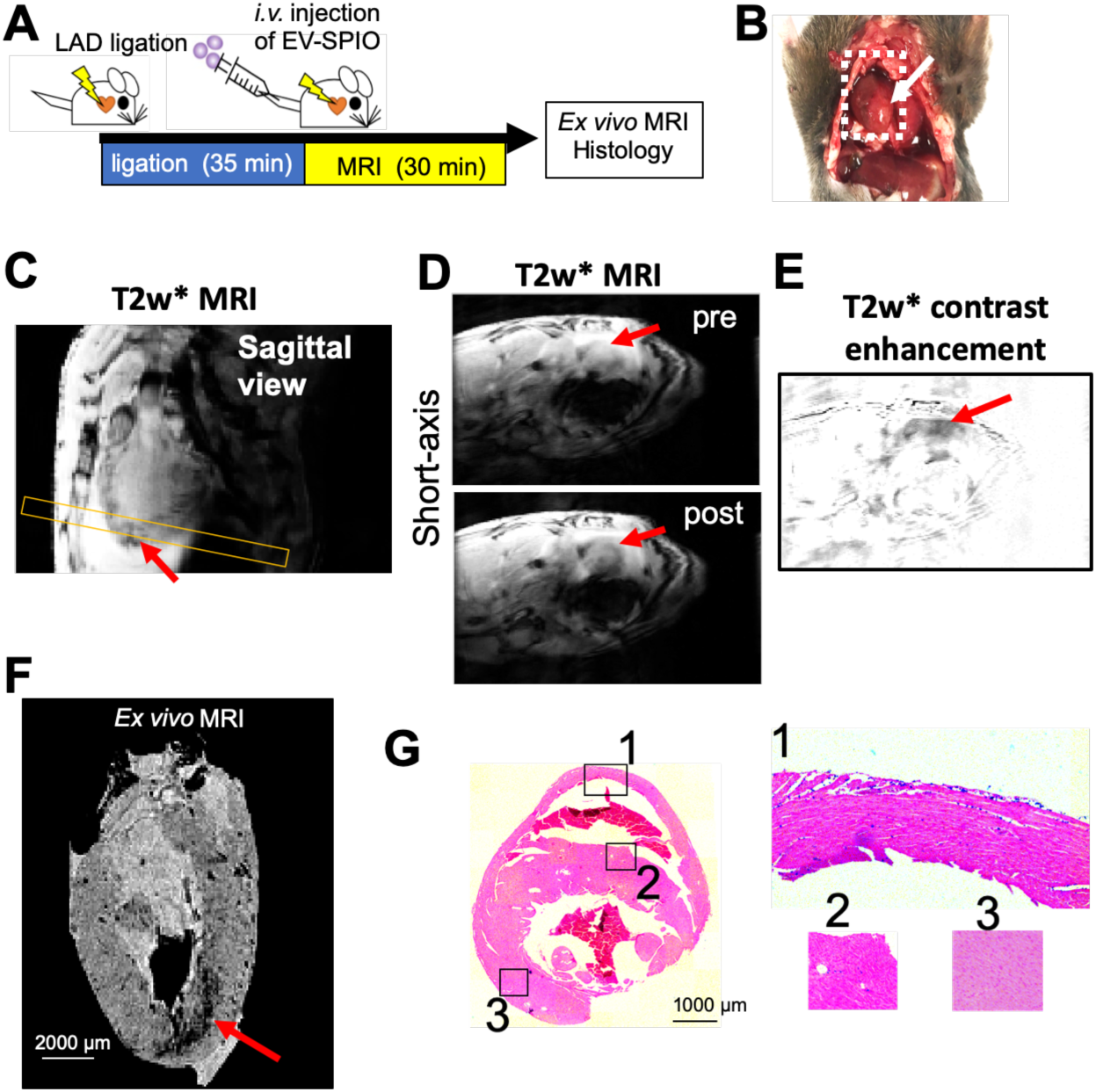
MRI tracking of magneto-EV accumulation in the IR heart. **A**. Schematic illustration of the experimental IR heart model and MRI acquisition. **B**. Macrophotograph of the heart with the IR region (arrow). **C**. Sagittal *in vivo* MR images of the heart. Yellow box indicates the slice position of the short-axis view. Short-axis pre- and post-injection *in vivo* T_2_*w images (**D**) and enhancement maps, defined as ΔT_2_*w=T_2_*w (post)-T_2_*w (pre) (**E**) showing hypointense areas in the injured region around the apex of the heart. **F**. *Ex vivo* heart MR image showing accumulation of magneto-EVs (red arrow). **G.** Prussian blue staining of the injured heart (Left: whole heart; Right: zoom-in of sections 1-3).

## Discussion

The primary aim of our study was to develop a non-invasive MRI tracking method for sensitive and accurate determination of the pharmacokinetics and biodistribution of therapeutic EVs in living subjects. Increasing evidence now supports the use of stem cell-derived EVs for regeneration of injured tissue with similar or even better therapeutic effects as compared to their parental cells, and without some of the limitations and risks associated with whole cell therapy-based approaches(*13, 38*). However, in order to optimize the use of EVs as therapeutic agents, much needs to be learned about their pharmacokinetics and whole-body distribution as related to their dose, injection route, and cell origin. To this end, the availability of a whole-body non-invasive imaging tracking method is critical. Indeed, for whole cell therapy, MRI has shown to be invaluable to monitor the spatiotemporal fate of injected cells in numerous preclinical studies and a few clinical studies(*39-41*). Our current study demonstrates that human iPSC-derived magneto-EVs produce a sufficiently strong MRI signal to allow for their *in vivo* tracking in the injured kidney and heart, in several different animal injury models.

One of the most challenging technical hurdles for imaging EVs is their small size. The typical diameter of EVs is less than 200 nm, making their volume approximately five orders of magnitude (1.25×10^5^ times) smaller than that of their parental cells (diameter ∼10 µm). We used SPIO nanoparticles as they have been shown to be the most sensitive MRI contrast agent(*42*). In addition, we used the smallest commercially available SPIO nanoparticles with a 5 nm core size (∼ 46 nm hydrodynamic size) to maximize the encapsulation rate. The TEM results showed that a few SPIO nanoparticles were loaded into a single EV, increasing detection sensitivity. The *in vivo* studies showed that a relatively low dose of 1.5 μg iron per mouse (0.67 μmol Fe/kg body weight) resulted in high MRI contrast allowing visualization of the accumulation of *i.v.*-injected magneto-EVs in injured tissue. In comparison, as an imaging agent for evaluation of renal function, SPIO has been used at a dose level of 60 µmol Fe/kg(*43*), two orders of magnitude higher than the dose used in the present study. Indeed, the estimated *in vivo* detection limit is only 8.76×10^7^ EVs/mL by our *in vitro* data. With such a high detectability, we were able to assess the biodistribution of iPSC-derived EVs using a low level of dose (*i.e.*, 1×10^9^ EVs or 10 µg proteins per mouse, assuming the protein content is 1 µg protein per 1×10^8^ EVs (*44*)), which is lower than those used for SPECT (*i.e.*, 29–64 µg (*14, 17*)), optical imaging (*i.e.*, 2.5-200 μg (*7, 14*)), and CT (*i.e.*, 28 μg (*19*)). To the best of our knowledge, there is no MRI studies reported using *i.v.* injection. While a report showed that local injection of 5×10^8^ EVs could result in a detectable MRI signal (*21*), higher doses have to be used to achieve a reliable MRI signal, *i.e.*, 25 µg in the same study and 50 µg for lymph node imaging with EVs injected in foot pads *(22).* Compared to all those pervious imaging methods, our technology provides a high sensitivity that permits MRI tracking of EVs *in vivo*.

Our platform technology for preparing highly purified magnetically labeled EVs is simple but efficient, and potentially useful for other nanoparticles and ligand-binding systems. Previous studies used commercially available SPIO nanoparticles by either pre-incubation of the parental cells with SPIOs, followed by the collection of labeled EVs(*21*), or ii) direct labeling with SPIOs using electroporation(*22, 45*). The first approach often suffers from low labeling efficiency as only a portion of EVs secreted by the labeled cells containing SPIOs, whereas the second approach relies on the use of sophisticated and time consuming (*i.e.*, 2 hours(*45*)) purification steps to remove unencapsulated SPIO. In contrast, our approach employs surface-modified SPIOs and the corresponding purification procedure is relatively simple, less equipment-intensive, and faster (∼minutes), providing an efficient way to prepare highly purified magneto-EVs with minimal EV loss during the purification steps. The resulting magneto-EVs have a high purity, hence allowing accurate detection and measurement of the spatiotemporal quantity and distribution of administrated EVs, without false positives (*i.e.*, non-encapsulated SPIOs).

Using the MRI signal of the magneto-EVs, we were able to detect the dynamic uptake and biodistribution of EVs in different tissues of three disease models non-invasively. In the LPS-AKI mice, the MRI results showed that magneto-EVs accumulated in the injured kidney at a much higher level than in uninjured controls. The distribution of EVs colocalized well with the injured proximal tubules in the cortex similar as previously reported for MSC-derived EVs in a cisplatin-induced AKI model(*46*). The accumulation of EVs in the injured areas was further confirmed in an IRI kidney model, where iPSC-derived EVs preferentially accumulated in damaged glomeruli in the medullar regions, in good agreement with previous reports on MSCs(*28, 47*). In addition, the accumulation of EVs provided significant protective effects and increased survival in the lethal LPS-AKI model. Moreover, our MRI results revealed similar distribution patterns for EVs isolated from iPSCs and FBS, suggesting that accumulation of EVs itself in the injured sites may not be attributable to the cell origin of EVs. However, only iPSC-derived EVs but not FBS-EVs produced a strong protective effect and improved the survival of mice with injured kidneys, supporting the notion that the therapeutic effects are attributed to the stem cell origin. Finally, unlike other synthetic nanoparticles, our MRI results revealed only a moderate uptake of EVs in the liver, even at a lower level than in the injured kidney, suggesting that EVs may be a promising new nanoparticulate drug carrier to bypass non-specific liver uptake and consequently improve drug delivery.

## Conclusions

We developed a platform technology for preparing magneto-EVs for *in vivo* MRI tracking. A high efficiency of purified magnetically labeled EVs could be achieved using SPIO particles functionalized with His-tags, while non-encapsulated particles could be easily removed using a Ni-NTA column. This strategy allowed successful preparation of magneto-EVs derived from either human iPSC or FBS. Highly purified labeled EVs are essential to mitigate false positives (*i.e.*, non-encapsulated SPIOs) for accurate detection and measurement of their spatiotemporal distribution. Magneto-EVs produced sufficient MRI signal to allow MRI detection of injected EVs in injured organs, including kidney and heart, correlating well with the histological assessment and therapeutic outcome in the LPS-AKI model. Our technology represents a novel tool for *in vivo* tracking of therapeutic EVs and for guiding their applications in EV-based regenerative medicine and possibly EV-based drug delivery.

## Materials and Methods

### Materials

Carboxyl SPIOs (SPIO-COOH, core diameter = 5 nm) was purchased from Ocean Nanotech (Springdale, AR). Hexa-histidine peptide was purchased from GeneScript (Springdale, AR, USA). 1-Ethyl-3-(3-dimethylaminopropyl)-carbodiimide (EDC), sulfo-N-hydroxysuccinimide (sulfo-NHS) and Ni-NTA were purchased from Sigma-Aldrich (St. Louis, MO, USA).

### Cell culture

Human iPSCs were programmed from blood cells of a healthy male donor(*48*), cultured in 6-well plates coated with vitronectin (Gibco, Calsbad, CA), and maintained in Essential 8 (E8) medium (Gibco) supplemented with 10 μM Y-27632 dihydrochloride (Stem Cell Technologies, Vancouver, Canada at 37°C in a humidified atmosphere of 5% CO_2_ and 95% air. The culture medium was changed daily after gently washing the cells with 10 mM PBS, pH=7.4. Cells were passaged at 80%-90% confluence using TrypLE™ Express Enzyme (Gibco) supplemented with 10 μM Y-27632 dihydrochloride. Cells were routinely checked for mycoplasma contamination.

### Synthesis of SPIO-His

Fifty microliters of SPIO-carboxyl (2 mg/mL) was mixed with 100 μg EDC (10 mg/mL) and 100 μg NHS (10 mg/mL), and 100 μL MES buffer (pH=6.0). The solution was shaken for 30 min. One mg of peptide (25 mg/mL in dd H_2_O) was then added, followed by adding 200 μL 10 mM PBS, pH=7.4. The pH of the solution was then adjusted to 7.4, followed by shaking for 2 h. The product was dialyzed, lyophilized and reconstituted to 2 mg Fe/mL.

### Collection, purification and enrichment of EVs

To acquire concentrated EVs, medium was harvested from multiple passages of iPSC culture and concentrated. Culture medium in 50 mL tubes was centrifuged for 10 min at 300 g followed by another 10 min at 2,000 g at 4 °C to remove cells and debris. The medium was then concentrated to 200 μL using an Amicon ultra-15 filter column and an Ultracel-100 membrane by centrifugation at 4,000 g for 20 min (MilliporeSigma, Billerica, MA, USA). Then, 0.5 mL of the concentrated medium was loaded to a qEV column (iZON, Cambridge, MA, USA) and eluted with PBS in 500 μL fractions with collection of the 7-9 fractions. This procedure removes small medium molecules from the concentrate. Purified EVs (about 2.4 mL elute from a total of 300 mL medium) were further concentrated six times using a qEV column to a final volume of 0.4 mL. To further enrich magneto-EVs, the microcentrifuge tube containing purified magneto-EVs was fixed upright on a 1-inch cube Neodymium magnet (CMS magnetics, Garland, TX, USA) overnight and the pelleted magneto-EVs were resuspended in the desirable volume of PBS. Isolation and purification of FBS (catalog #F2442, Sigma-Aldrich) derived EVs were performed in a similar manner.

### Electroporation of EVs

Fifty microliters of concentrated EVs (1.1×10^11^/mL) were mixed with 25 μL of 2 mg/mL SPIO-His. Electroporation was performed with a Gene Pulser Xcell Electroporation Systems (Biorad) using two pulses of 240 V/mm and 100 F Capacitance with a 1 mm cuvette. EVs were then transferred to a clean microcentrifuge tube and placed on ice for 1 h before Ni-NTA purification.

### Purification of magneto-EVs

Ni-NTA columns were prepared by packing 1 mL Ni-NTA His•Bind resins (Sigma) into a 6-ml ISOLUTE® Single Fritted Reservoir column with 10 μm polyethylene frit (Biotage, Charlotte, NC, USA), followed by washing with 5 mL PBS. After electroporation, EVs (∼75 μL) were resuspended in 200 μL PBS and then loaded onto a Ni-NTA column and gently shaken for 30 min. After the first elute was collected, the column was rinsed with 200 μL PBS. All elutes were collected using 1 mL microcentrifuge tubes.

The size and iron content of EVs was measured by dynamic light scattering (DLS, Nanosizer ZS90, Malvern Instruments) and by inductively coupled plasma optical emission spectroscopy (ICP-OES, Thermo iCAP 7600, ThermoFisher Scientific, Waltham, MA, USA), respectively. To estimate the loss of EVs during the purification procedure, the numbers of EVs before and after purification were measured using a nanoparticle tracking analysis (NTA) instrument (Zetaview, Particle Metrix, Germany) using a 488-nm laser and ZetaView 8.04.02 software.

### Transmission electron tomography

Carbon-coated 400 mesh copper grids (Electron Microscopy Services, Hatfield, PA, USA) were placed on 30 μl drops of SPIO or EV samples for one minute. Grids were quickly washed with two drops of dH_2_O and blotted dry on a filter paper. Grids were stained for one minute with 2% uranyl acetate (Electron Microscopy Services) in dH_2_0 and blotted dry. All imaging was performed with a Zeiss Libra 120 TEM operated at 120 KV and equipped with an Olympus Veleta camera (Olympus Soft Imaging Solutions GmbH, Münster, Germany).

### Mouse models of acute kidney injury

All animal experiments were approved by our Animal Care and Use Committee. Male C57BL/6J mice (6-8 weeks of age) were acquired from Jackson Laboratories (Bar Harbor, ME, USA). The LPS-AKI model was established as described previously(*29*) by intraperitoneal (*i.p.*) administration of lipopolysaccharide (LPS, Sigma Aldrich) at a dose of 10 mg/kg. The IRI-AKI model was established according to a previously published procedure(*49*). In brief, animals were anesthetized with 2% isoflurane and an incision was made in the back muscle and skin layer to expose the right kidney and the renal vascular pedicle was clamped using a microvessel clamp (#18052-03, Fine Science Tools, Foster City, CA, USA) for 45 min, followed by suture-closing the incisions.

### Mouse model of heart ischemic and reperfusion injury (IRI)

Male C57BL/6J mice under induction anesthesia with 3-4% isoflurane received 0.03-0.07 mg/kg buprenorphine subcutaneously (*s.c.*). Isoflurane was then adjusted to 1-2% and 2 mg/kg succinylcholine was administered *i.p.* After 5 min, a left thoracotomy was performed in the 5th to 6th intercostal space. The pericardium was torn and the coronary artery was located. A piece of 7-0 prolene was tied around the coronary artery with a small piece of black polyethylene PE10 tubing (Braintree Scientific Inc., Braintree, MA, USA) under the suture. Occlusion was performed for 35 min. During occlusion, ribs were closed with one single 5-0 silk suture and the skin was closed with a bulldog clamp (Fine Science Tools, Foster City, CA, USA). At 5 min prior to ending the occlusion, the chest was reopened carefully and the sutures were removed. Ribs and skin were closed with 5-0 silk. Mice were allowed to regain consciousness and a second dose of buprenorphine at the dose of 0.06-0.075 mg/kg was administered *s.c*.

### In vitro MRI

Samples of SPIO-COOH, SPIO-His, unlabeled and labeled EVs were prepared at different concentrations in 10 mM PBS, pH=7.4 and transferred to 5 mm glass NMR tubes, and then combined for MRI measurements on a Bruker 9.4 Tesla vertical bore scanner equipped with a 20 mm birdcage transmit/receive coil. T_2_ relaxation times were measured using the Carr-Purcell-Meiboom-Gill (CPMG) method at room temperature as previously described(*50*). The acquisition parameters were: TR/TE=25 s/4.3 ms, RARE factor=16, matrix size=64×64, in-plane resolution=0.25×0.25 mm and slice thickness=2 mm. Each T_2_w image took approximately 1’40’’ to acquire. The T_2_ relaxivities (r_2_) were calculated based on mean R_2_ (=1/T_2_) of each sample and their concentrations (c), using the following equation:

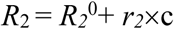

Where R_2_^0^ represents the inherent water proton transverse relaxation rate.

### In vivo MRI

All animal studies were performed on a 11.7T Biospec (Bruker) horizontal bore scanner equipped with a mouse brain surface array RF coil (receiver) and a 72 mm volume coil (transmitter). A 30-min dynamic scan was acquired using a fast low angle shot (FLASH) gradient echo sequence immediately before *i.v.* injection of 1×10^9^ magneto-EVs or 1.5 μg SPIO-His (having the same iron amount as that in magneto-EVs) in 200 μL PBS. The acquisition parameters were: flip angle=25°, TR=800 ms, TE=5.8 ms, matrix size=256×128 and resolution=167×280 mm^2^. Before and 30 min after injection T_2_* maps were also acquired using a multiple gradient echo (MGE) pulse sequence with TR=800 ms and TE times of 2.6, 5.8, 9, 12.2, 15.4, 18.6, 21.8, and 25 ms.

After MRI, mice were euthanized by cervical dislocation under anesthesia and tissues of interest were collected and fixed in 4% paraformaldehyde solution for *ex vivo* MRI and histological analysis.

### Ex vivo MRI

Excised organs (*i.e.*, kidney, heart, and liver) were transferred to a 5 ml syringe filled with proton-free fluid Fomblin (Solvay Solexis, Inc., USA) and *ex vivo* high-resolution MRI was performed on a vertical bore 9.4T Bruker scanner equipped with a 15 mm birdcage transmit/receive volume coil. A three-dimensional FLASH sequence was used with TE=6 ms, TR=150 ms, matrix size=310×230×155, FOV=12×9×6 mm, resolution=0.039×0.039×0.039 mm, averages=5, and flip angle=15°. The total scan time was 6h43m12s.Amira 3D Visualization Software 5.4.3 (Visage Imaging Inc., Carlsbad, CA, USA) was used to quantify the areas of hypointense signal and to visualize the 3D distribution of magneto-EVs.

### Histological analysis

Excised tissues were paraffin-embedded and sectioned at 5 μm thickness, and stained with Prussian blue and Periodic acid–Schiff (PAS). Sections were imaged using a Zeiss Axio Observer Z1 microscope (Zeiss, Oberkochen, Germany) and processed using Zen Pro software.

### Effects of iPSC-EV treatment on LPS-induced AKI mice

Five treatment groups were tested: 1) iPSC-EV was administered at the same time as LPS injection; 2) iPSC-EV was administered at 3 hours after LPS injection; 3) iPSC-EV was administered at 24 hours after LPS injection; 4) FSB-EV was administered at 3 hours after LPS injection; and 5) vehicle control (200 μL PBS) was administered at the same time as LPS injection. Five mice were randomly chosen for each group. For all EV treatment groups, 2×10^9^ EVs in 200 μL PBS were administered *i.v.* through the tail vein. Animal survival was monitored for each group every day up to 6 days.

### Statistical analysis

All data are presented as mean±s.e.m. GraphPad Prism version 8 (GraphPad Software Inc., San Diego, CA, USA) was used to perform statistical analysis. A unpaired two-tailed Student’s t-test was used to compare the difference between two groups. Differences with *P*<0.05 were considered statistically significant. The Kaplan-Meier method was used to analyze animal survival data.

## Supporting information

### Other Supplementary files

**Video 1**. Dynamic T_2_*w images of a normal mouse after injection of magneto-EVs.

**Video 2**. Dynamic T_2_*w images of an LPS-AKI mouse after injection of magneto-Evs

**Video 3**. 3D visualization of the biodistribution of magneto-EVs in a LPS-AKI kidney.

**Figure S1.**
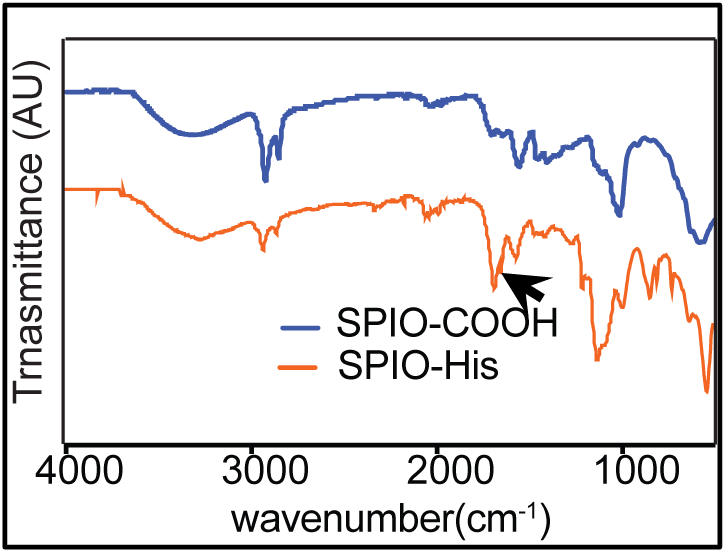
Fourier-transform infrared spectrum of SPIO-His compared to that of SPIO-COOH which shows the characteristic peak of SPIO-His near 1680 cm^-1^, confirming the formation of amide bonds between carboxyl group and the amine group of histidine peptide.

**Figure S2.**
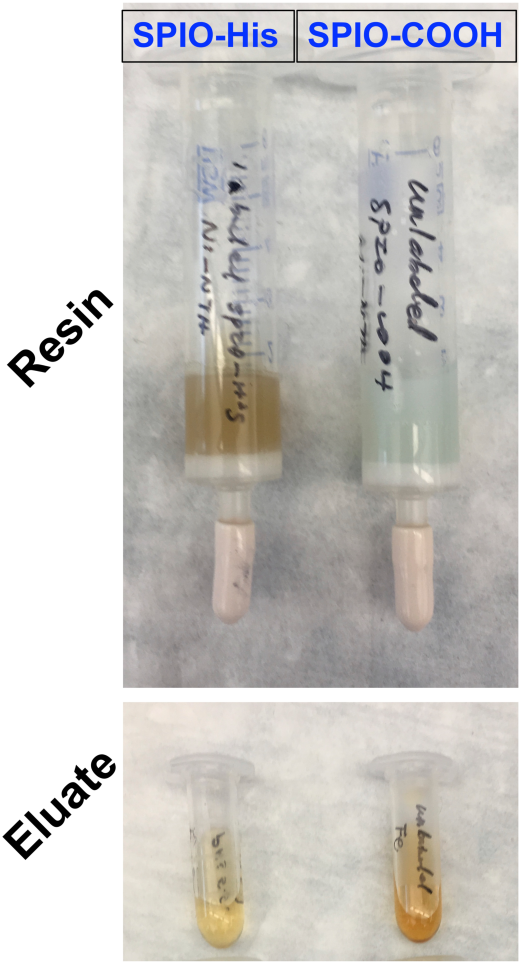
Photos of the Ni-NTA column and eluates after washing with SPIO solutions. In brief, equivalent amounts (0.1 mg) of SPIO-His and SPIO-COOH were passed through columns packaged with Ni-NTA resin. Then, the columns were washed using PBS and all eluates were collected. The Ni-NTA columns absorbing SPIO-His appeared brownish, whereas the columns passed with SPIO-COOH remained light blue; the color of SPIO-His eluate was much lighter than that of SPIO-His.

**Figure S3.**
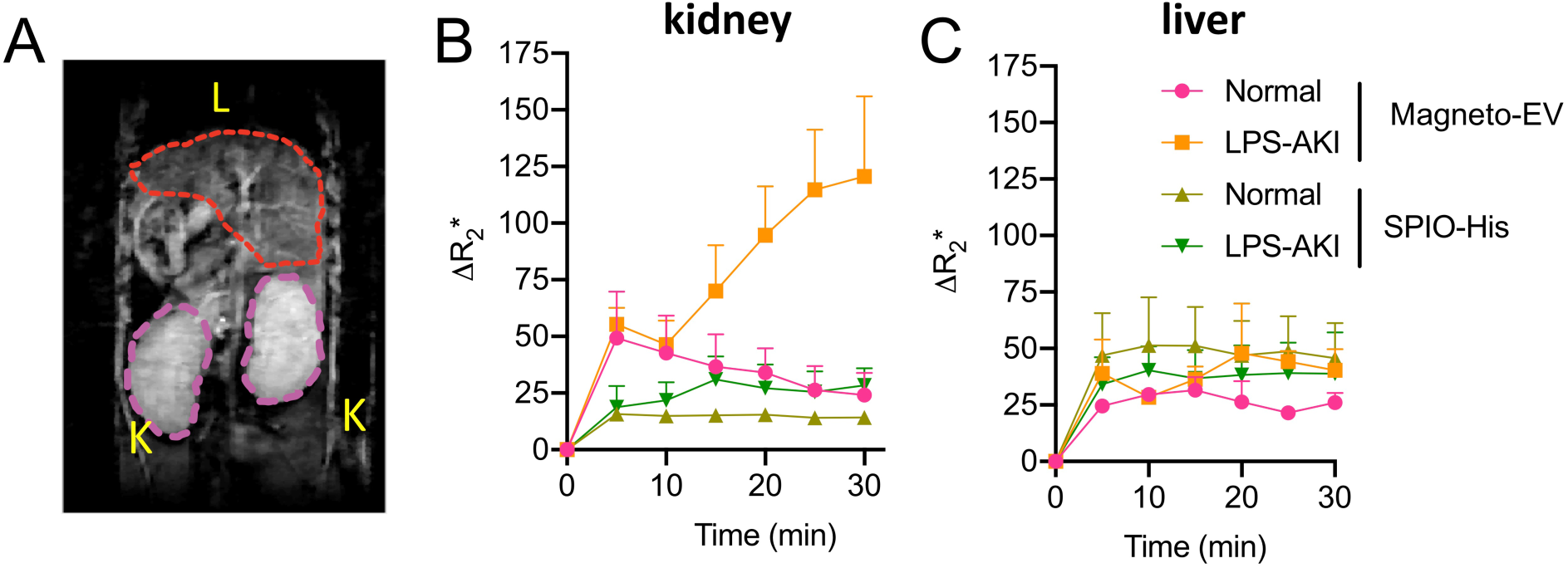
Dynamics of ΔR_2_* contrast change in kidney and liver generated by magneto-EVs. **A.** Coronal T_2_*w image of a representative mouse showing the ROI selection for kidneys and liver. Dynamic changes of ΔR_2_*, calculated by 1/T_E_*×ln(S^post^/S^pre^), of kidney (**B**) and liver (**C**) of normal and LPS-AKI mice after magneto-EV or SPIO-His injection.

**Figure S4.**
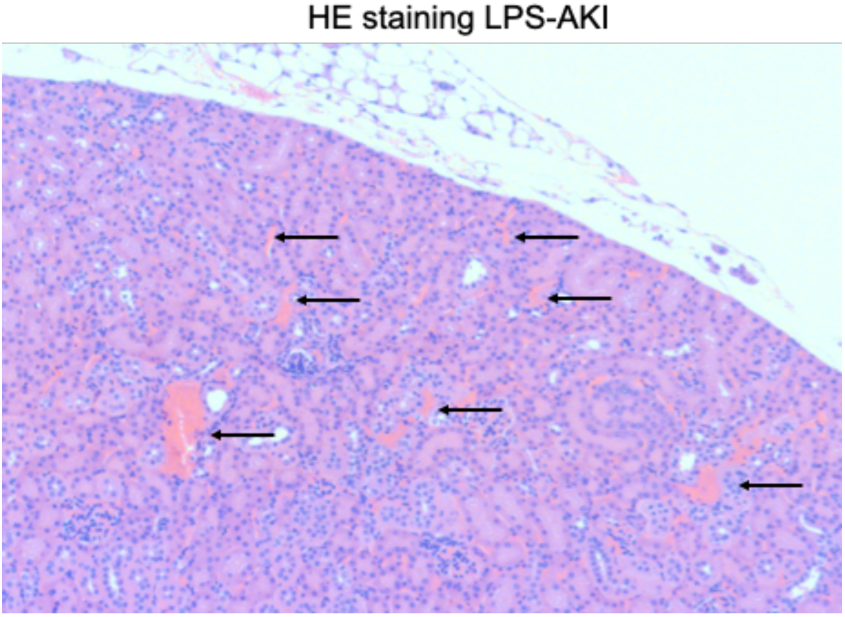
Hematoxylin-eosin (HE) stain of a representative section of LPS-AKI kidney in which hemorrhages in the cortex of the kidney are indicated by black arrows.

## References

1. D. Karpman, A. L. Stahl, I. Arvidsson, Extracellular vesicles in renal disease. Nature reviews. Nephrology 13, 545 (Sep, 2017).

2. G. van Niel, G. D’Angelo, G. Raposo, Shedding light on the cell biology of extracellular vesicles. Nat Rev Mol Cell Biol 19, 213 (Apr, 2018).

3. M. Colombo, G. Raposo, C. Thery, Biogenesis, secretion, and intercellular interactions of exosomes and other extracellular vesicles. Annu Rev Cell Dev Bi 30, 255 (2014).

4. M. Gnecchi, Z. Zhang, A. Ni, V. J. Dzau, Paracrine mechanisms in adult stem cell signaling and therapy. Circ Res 103, 1204 (Nov 21, 2008).

5. L. Chen, E. E. Tredget, P. Y. Wu, Y. Wu, Paracrine factors of mesenchymal stem cells recruit macrophages and endothelial lineage cells and enhance wound healing. PLoS One 3, e1886 (Apr 2, 2008).

6. S. Rani, A. E. Ryan, M. D. Griffin, T. Ritter, Mesenchymal Stem Cell-derived Extracellular Vesicles: Toward Cell-free Therapeutic Applications. Mol Ther 23, 812 (May, 2015).

7. I. Vishnubhatla, R. Corteling, L. Stevanato, C. Hicks, J. Sinden, The Development of Stem Cell-Derived Exosomes as a Cell-Free Regenerative Medicine. Journal of Circulating Biomarkers 3, (2014).

8. F. Collino et al., AKI Recovery Induced by Mesenchymal Stromal Cell-Derived Extracellular Vesicles Carrying MicroRNAs. Journal of the American Society of Nephrology : JASN 26, 2349 (Oct, 2015).

9. L. A. Reis et al., Bone marrow-derived mesenchymal stem cells repaired but did not prevent gentamicin-induced acute kidney injury through paracrine effects in rats. PLoS One 7, e44092 (2012).

10. S. Bruno et al., Mesenchymal stem cell-derived microvesicles protect against acute tubular injury. Journal of the American Society of Nephrology : JASN 20, 1053 (May, 2009).

11. S. Bruno et al., Microvesicles derived from mesenchymal stem cells enhance survival in a lethal model of acute kidney injury. PLoS One 7, e33115 (Mar 14, 2012).

12. M. Adamiak et al., Induced Pluripotent Stem Cell (iPSC)-Derived Extracellular Vesicles Are Safer and More Effective for Cardiac Repair Than iPSCs. Circ Res 122, 296 (Jan 19, 2018).

13. B. Liu et al., Cardiac recovery via extended cell-free delivery of extracellular vesicles secreted by cardiomyocytes derived from induced pluripotent stem cells. Nat Biomed Eng 2, 293 (May, 2018).

14. C. Grange et al., Biodistribution of mesenchymal stem cell-derived extracellular vesicles in a model of acute kidney injury monitored by optical imaging. International journal of molecular medicine 33, 1055 (May, 2014).

15. O. P. Wiklander et al., Extracellular vesicle in vivo biodistribution is determined by cell source, route of administration and targeting. J Extracell Vesicles 4, 26316 (2015).

16. T. Imai et al., Macrophage-dependent clearance of systemically administered B16BL6-derived exosomes from the blood circulation in mice. J Extracell Vesicles 4, 26238 (2015).

17. D. W. Hwang et al., Noninvasive imaging of radiolabeled exosome-mimetic nanovesicle using (99m)Tc-HMPAO. Sci Rep 5, 15636 (Oct 26, 2015).

18. S. Shi et al., Copper-64 Labeled PEGylated Exosomes for In Vivo Positron Emission Tomography and Enhanced Tumor Retention. Bioconjug Chem 30, 2675 (Oct 16, 2019).

19. O. Betzer, N. Perets, E. Barnoy, D. Offen, R. Popovtzer, Labeling and tracking exosomes within the brain using gold nanoparticles. Proc Spie 10506, (2018).

20. N. Perets et al., Golden Exosomes Selectively Target Brain Pathologies in Neurodegenerative and Neurodevelopmental Disorders. Nano Lett 19, 3422 (Jun 12, 2019).

21. A. Busato et al., Magnetic resonance imaging of ultrasmall superparamagnetic iron oxide-labeled exosomes from stem cells: a new method to obtain labeled exosomes. International journal of nanomedicine 11, 2481 (2016).

22. L. Hu, S. A. Wickline, J. L. Hood, Magnetic resonance imaging of melanoma exosomes in lymph nodes. Magnetic resonance in medicine 74, 266 (Jul, 2015).

23. K. O. Jung, H. Jo, J. H. Yu, S. S. Gambhir, G. Pratx, Development and MPI tracking of novel hypoxia-targeted theranostic exosomes. Biomaterials 177, 139 (Sep, 2018).

24. P. Y. Lee et al., Induced pluripotent stem cells without c-Myc attenuate acute kidney injury via downregulating the signaling of oxidative stress and inflammation in ischemia-reperfusion rats. Cell Transplant 21, 2569 (2012).

25. S. Liu et al., Highly Purified Human Extracellular Vesicles Produced by Stem Cells Alleviate Aging Cellular Phenotypes of Senescent Human Cells. Stem Cells 37, 779 (Jun, 2019).

26. J. H. Jung, X. Fu, P. C. Yang, Exosomes Generated From iPSC-Derivatives: New Direction for Stem Cell Therapy in Human Heart Diseases. Circ Res 120, 407 (Jan 20, 2017).

27. Y. Shi, H. Inoue, J. C. Wu, S. Yamanaka, Induced pluripotent stem cell technology: a decade of progress. Nat Rev Drug Discov 16, 115 (Feb, 2017).

28. H. Ittrich et al., In vivo magnetic resonance imaging of iron oxide-labeled, arterially-injected mesenchymal stem cells in kidneys of rats with acute ischemic kidney injury: detection and monitoring at 3T. J Magn Reson Imaging 25, 1179 (Jun, 2007).

29. J. Liu et al., CEST MRI of sepsis-induced acute kidney injury. NMR Biomed 31, e3942 (Aug, 2018).

30. K. Doi, A. Leelahavanichkul, P. S. Yuen, R. A. Star, Animal models of sepsis and sepsis-induced kidney injury. J Clin Invest 119, 2868 (Oct, 2009).

31. J. W. Bulte, I. D. Duncan, J. A. Frank, In vivo magnetic resonance tracking of magnetically labeled cells after transplantation. J Cereb Blood Flow Metab 22, 899 (Aug, 2002).

32. J. Huang, X. Zhong, L. Wang, L. Yang, H. Mao, Improving the magnetic resonance imaging contrast and detection methods with engineered magnetic nanoparticles. Theranostics 2, 86 (2012).

33. M. Morishita, Y. Takahashi, M. Nishikawa, Y. Takakura, Pharmacokinetics of Exosomes-An Important Factor for Elucidating the Biological Roles of Exosomes and for the Development of Exosome-Based Therapeutics. J Pharm Sci 106, 2265 (Sep, 2017).

34. Y. Feng et al., Improved MRI R2 * relaxometry of iron-loaded liver with noise correction. Magnetic resonance in medicine 70, 1765 (Dec, 2013).

35. P. Reimer, B. Tombach, Hepatic MRI with SPIO: detection and characterization of focal liver lesions. Eur Radiol 8, 1198 (1998).

36. L. Zhang et al., Nerolidol Protects Against LPS-induced Acute Kidney Injury via Inhibiting TLR4/NF-kappaB Signaling. Phytother Res 31, 459 (Mar, 2017).

37. J. M. J. Pickard, N. Burke, S. M. Davidson, D. M. Yellon, Intrinsic cardiac ganglia and acetylcholine are important in the mechanism of ischaemic preconditioning. Basic Res Cardiol 112, 11 (Mar, 2017).

38. S. R. Baglio, D. M. Pegtel, N. Baldini, Mesenchymal stem cell secreted vesicles provide novel opportunities in (stem) cell-free therapy. Front Physiol 3, 359 (2012).

39. J. W. Bulte, In vivo MRI cell tracking: clinical studies. AJR Am J Roentgenol 193, 314 (Aug, 2009).

40. J. W. M. Bulte, H. E. Daldrup-Link, Clinical Tracking of Cell Transfer and Cell Transplantation: Trials and Tribulations. Radiology 289, 604 (Dec, 2018).

41. I. J. de Vries et al., Magnetic resonance tracking of dendritic cells in melanoma patients for monitoring of cellular therapy. Nat Biotechnol 23, 1407 (Nov, 2005).

42. J. W. Bulte, D. L. Kraitchman, Iron oxide MR contrast agents for molecular and cellular imaging. NMR Biomed 17, 484 (Nov, 2004).

43. J. P. Laissy et al., Reversibility of experimental acute renal failure in rats: assessment with USPIO-enhanced MR imaging. J Magn Reson Imaging 12, 278 (Aug, 2000).

44. F. Pi et al., Nanoparticle orientation to control RNA loading and ligand display on extracellular vesicles for cancer regression. Nat Nanotechnol 13, 82 (Jan, 2018).

45. J. L. Hood, M. J. Scott, S. A. Wickline, Maximizing exosome colloidal stability following electroporation. Anal Biochem 448, 41 (Mar 1, 2014).

46. Y. Zhou et al., Exosomes released by human umbilical cord mesenchymal stem cells protect against cisplatin-induced renal oxidative stress and apoptosis in vivo and in vitro. Stem Cell Res Ther 4, 34 (Apr 25, 2013).

47. R. Zhang, J. Li, L. Xin, J. Xie, In vivo magnetic resonance imaging of iron oxide-labeled, intravenous-injected mesenchymal stem cells in kidneys of rabbits with acute ischemic kidney injury: detection and monitoring at 1.5 T. Renal Failure 37, 1363 (2015).

48. B. K. Chou et al., Efficient human iPS cell derivation by a non-integrating plasmid from blood cells with unique epigenetic and gene expression signatures. Cell Res 21, 518 (Mar, 2011).

49. Q. Wei, Z. Dong, Mouse model of ischemic acute kidney injury: technical notes and tricks. Am J Physiol Renal Physiol 303, F1487 (Dec 1, 2012).

50. J. Zhang et al., Triazoles as T2 -Exchange Magnetic Resonance Imaging Contrast Agents for the Detection of Nitrilase Activity. Chemistry (Weinheim an der Bergstrasse, Germany) 24, 15013 (Oct 9, 2018).

